# Combined negative effects of simultaneous perturbations on grassland functionality

**DOI:** 10.64898/2026.09.28.754907

**Authors:** Marco Fioratti Junod, Sophie Gombeer, Julia Holmes, Stephan Zimmerman, Matthias C. Rillig, Anita Christina Risch, Irene Cordero

## Abstract

1. Grassland ecosystems are pivotal for global biodiversity, carbon (C) storage and food security. This biome, however, together with the ecosystem services it provides, is threatened by global change perturbations. These perturbations rarely occur in isolation and most often affect environments simultaneously, which is a pressing issue in ecology. Nevertheless, most of the research conducted so far has focussed on single simultaneous perturbations or pairs thereof. The few examples considering the effect of more than two simultaneous perturbations focussed only on plants or soil separately, which ignores the potential buffering or feedback effects on ecosystem functioning induced by plant-soils interactions.
2. To tackle this knowledge gap, we used an experimental setup involving mesocosms (soil and plants intact) collected in an extensively managed permanent grassland. Mesocosms were arranged in a greenhouse experiment where we applied individual and combined global change perturbations (up to ten in total) including application of fertilisers, pesticides and antibiotics, simulated trampling and grazing, heat stress and drought. Measured ecosystem functions and properties included a large set of biological, physical and chemical soil and plant associated variables.
3. Results showed that multiple perturbations in most cases had negative effects irrespective of the direction found for individual perturbations. However, in contrast to previous bare soil experiments, we only found scarce evidence for substantial synergistic effects of simultaneous perturbations in our experiment conducted with mesocosms that contained vegetation. This suggests a mitigating impact of plants on soil responses to global change. Moreover, we found a strong dominative effect of drought on ecosystem functioning, driving most of the effects observed.
4. Synthesis: Our results highlight the importance of evaluating global change impacts under more realistic settings, i.e., applying multiple perturbations simultaneously and including plant-soil interactions. Mitigation of global change in terrestrial ecosystems will only be possible once we holistically understand how ecosystems respond to altered conditions.

## Introduction

Global change is one of the most pressing challenges of our time. Extreme weather events, such as drought and heatwaves, are increasing rapidly in intensity and frequency (Fischer et al., 2021; IPCC, 2021), severely impacting terrestrial ecosystems (Bellard et al., 2012; Malhi et al., 2020). At the same time land use intensification is the worldwide norm to respond to the resource needs of an ever-growing human population (Pellegrini & Fernández, 2018). This intensification includes increasing use of fertilisers and pesticides (i.e., herbicides, fungicides, insecticides) and higher livestock densities, which leads to higher rates of defoliation and trampling. Together, these different global change impacts, hereafter perturbations, have far-reaching and often negative consequences for the environment, plant and animal biodiversity, food security, and ultimately human well-being (Steffen et al., 2002).

Grassland ecosystems cover ∼40% of the terrestrial area (O’Mara, 2012), store 20-30% of the terrestrial carbon (C) (Dobson et al., 2022; Eswaran et al., 1993), are a global reservoir of biodiversity (Petermann & Buzhdygan, 2021), and are of major economic and ecological importance (Gibson, 2009). In these ecosystems many different perturbations act simultaneously, e.g., grazing, trampling, fertilisation, or pesticide additions and livestock excretes containing considerable amounts of the antibiotics they were treated with (Kuppusamy et al., 2018), which together could impact food security and jeopardise climate change mitigation (O’Mara, 2012). Therefore, it is of paramount importance to understand grassland functional responses to multiple simultaneous global change perturbations. But assessing the effects of simultaneous perturbations is not straightforward. It has been shown, for example, that different individual perturbations can have opposing effects on ecosystems functioning: fertilisation enhanced soil microbial biomass (C. Chen & Xiao, 2023), while the use of pesticides reduced it (Pathak et al., 2022; Wołejko et al., 2020). Hence, is has to be expected that these two perturbations would compensate each other when they occurred simultaneously and result in a broadly net neutral effect in the response variable of interest. However, recent research demonstrated that different global change perturbations occurring simultaneously can have directional negative effects, even when individually some showed positive effects (Rillig et al., 2019).

According to multiple-stressor theory (Côté et al., 2016), simultaneous perturbations can produce four outcomes: i) Additive effects when their combined effect equals the algebraic sum of individual effects. ii) Dominative effects if the resulting effect is the same as the one of the strongest perturbation applied individually. iii) Antagonistic effects when the combined effect is less than the sum of each individual perturbation, and therefore less than expected additively. An example of an antagonistic response is the proportional model, where the individual effects are divided by the total number of perturbations applied before being added up. iv) Synergistic effects when the combined effect is greater than the sum of each perturbation independently, and therefore the combined response exceeds the additive expectation. An example of a synergistic response is represented by the augmentative model, where each individual effect is multiplied by a factor greater than 1 (Fig. 1) (Côté et al., 2016).

**Fig. 1:**
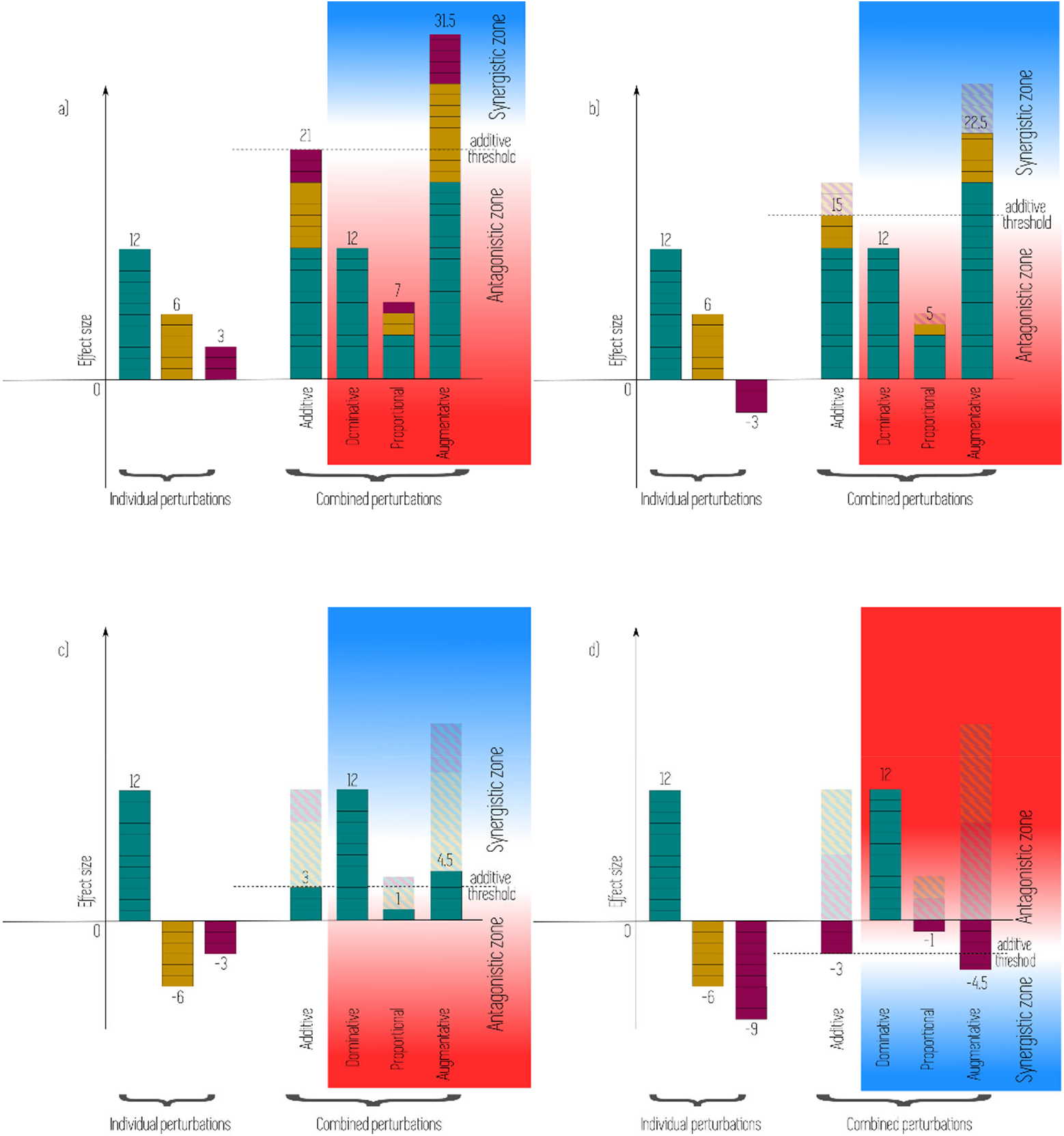
Conceptual figure of the potential responses of ecosystem properties when three individual perturbations are applied in combination. The effect size, i.e. difference to the control, is shown. In all cases, we show an additive model (algebraic addition of individual effects), which sets the threshold between the antagonistic and the synergistic response zone. Additionally, we show the dominative model (response equals the strongest individual effect), a proportional model (individual effects are divided by the number of perturbations applied) and an augmentative model (an arbitrary example of a synergistic response where each individual effect is multiplied by 1.5). Dashed areas represent the subtraction of the effects. a) All three individual perturbations have a positive effect on the response variable. b) One of the individual perturbations has a negative effect on the response variable. c) An example where the dominative model falls within the synergistic zone. d) The additive model predicts a decline in the response variable and therefore the synergistic zone is in the negative region.

Yet, to date, the majority of research has only considered global change perturbations individually or in pairs and we lack an understanding of the response of ecosystems to multiple simultaneous perturbations, which is essential since in nature perturbations rarely occur in isolation (Rineau et al., 2019). Only a handful of studies combined three or four perturbations and showed highly contrasting results, some reporting additive effects (Gutknecht et al., 2010; Niboyet et al., 2011), while others found antagonistic ones (Henry et al., 2005; Hines et al., 2017). Recent research on multiple perturbations (up to ten different ones) showed, in contrast, synergistic effects of these perturbations on plant and soil functions and diversity (Bi et al., 2024; Meidl et al., 2024; Rillig et al., 2019; Speißer et al., 2022; Zandalinas et al., 2021). Similarly, some recent publications came to the conclusion that multiple simultaneous global change perturbations would affect soil functions in a synergistic way (Komatsu et al., 2019; Rillig et al., 2023; Yang et al., 2021). However, the few experimental studies outlined above either focussed on plants alone or were conducted as bare soil experiments that did not include plants (Bi et al., 2024; Meidl et al., 2024; Rillig et al., 2019). Thus, we lack evidence of how interactions between plants and soil will help mitigate different, simultaneous global change perturbations. These plant-soil interactions for example, include the effect of plants on soil communities through root exudates or litter inputs (Chomel et al., 2019; de Vries et al., 2019; Karlowsky et al., 2018; Leff et al., 2018; Wardle et al., 2004; Williams & de Vries, 2020). In addition, plants can take up added nutrients or pesticides (Hussain et al., 2009), which could lessen the negative responses of these inputs on soil chemical and biological properties. However, if perturbation effects are strong enough to lead to plant death, the mitigating effects of the plants on soil properties could be reverted.

In this study, we addressed this knowledge gap by assessing how multiple, simultaneous global change perturbations affect ecosystem functioning as well as plant and soil properties. We used intact grassland mesocosms collected in the field containing plants in their original soil for our experiment. This design and experimental setup allowed us to account for plant-soil interactions in ecosystem response to multiple, simultaneous global change perturbations. Based on previous literature (Bi et al., 2024; Meidl et al., 2024; Rillig et al., 2019; Speißer et al., 2022; Zandalinas et al., 2021), we hypothesised that several grassland functions will respond in a synergistic way to multiple combined perturbations. Therefore, we expect that soil functionality will show stronger reductions the more perturbations we apply. However, we also hypothesised that the presence of the plants will reduce the magnitude of these synergistic effects.

## Materials and methods

### Experimental design

Grassland mesocosms (15 cm diameter, 15 cm deep) including soil and natural vegetation, were collected by driving PVC pipes with a 5 mm thick wall (Supplementary Fig. 1) into the soil of an extensively managed grassland (no fertilisation, no ploughing, 2 cuts per year with removal of the biomass). The grassland is located on the grounds of the Swiss Federal Institute for Forest, Snow and Landscape Research (WSL), Switzerland (47° 21’ 43.6” N; 8° 27’ 27.4” E), and dominated by *Dactylis glomerata*, *Holcus lanatus* and *Lolium* sp. (full species list in Supplementary Table 1). Mesocosm studies have demonstrated to be an experimental approach closer to real field conditions than pot experiments, in which many environmental factors can be manipulated. Moreover, they allow the application of toxic pollutants not suitable for the field (Cueff et al., 2020; Oram et al., 2020; Osburn et al., 2021). The mesocosms were collected on the 8^th^ (batch 1) and 22^nd^ of June 2023 (batch 2), approximately two weeks after mowing the patches where the mesocosms were collected. The mesocosm collection was split into two batches for logistical reasons, mainly manpower. We added “batch” to our statistical models to account for heterogeneity between batches in our data analyses. We covered the bottom of the mesocosms with a water permeable cloth immediately after collection to avoid soil loss (Supplementary Fig. 1) and put them into a greenhouse with 23 °C average temperature and 67% average air humidity. After 3.5 weeks (25 days) of acclimation, experimental treatments started (see below). After 5 weeks of treatment application, we harvested the mesocosms. During the experiment, mesocosms were rotated once per week to avoid confounding effects due to their position in the greenhouse.

Ten different global change perturbations were applied randomly, individually and in combination, at an ecologically relevant level. *Fertiliser additions:* nitrogen (N): NH_4_NO_3_ equivalent of 50 kg ha^-1^ yr^-1^ of N and phosphorus (P): Ca(H_2_PO_4_)_2_ · H_2_O equivalent of 50 kg ha^-1^ yr^-1^ of P, as the recommended doses of fertiliser added to pastures in Switzerland for low to medium intensity management (Richner et al., 2017). *Defoliation*: repeated cutting of the aboveground plant biomass to a sward height of 4 cm, which is a widely recommend sward height for grazed grasslands (Medina-Roldán & Bardgett, 2011). We cut the vegetation once a week to simulate intensive grazing (Medina-Roldán & Bardgett, 2011). *Trampling*: once a week, to match the defoliation frequency, and simulating an adult cow. We dropped a 10 kg weight from 0.3 m height onto a cow claw with a 48 cm^2^ basal surface area. Through this the soil compacted approximately 2 cm on impact, resulting in a dynamic pressure of approximately 300 kPa, which is equivalent of an adult cow (Striker et al., 2006). *Insecticide*: thiamethoxam, 96 g ha^-1^, and *fungicide:* carbendazim 150 g ha^-1^, which are the recommended doses of the commercial products for pastures. *Herbicide*: glyphosate 1.64 kg ha^-1^, which is the average dose used in pastures in Switzerland (Keiser & Ramsebner, 2020). *Antibiotics:* 10 mg oxytetracycline, which will simulate a fresh deposition of manure (Kuppusamy et al., 2018)*. Drought*: reduction of water-holding capacity (WHC) from 60% to 30%. Soil moisture levels were controlled by weighing and watering the mesocosms three times per week. For the drought treatment, watering was withheld until the individual mesocosms reached the target soil moisture of 30% WHC. This drought level equals 0.14 volumetric soil water content, which corresponds to a drought frequently experienced in the region (return period of 1.45 years). The return period was calculated using the ESA CCI soil moisture database (Dorigo et al., 2023) cut to the lowlands of Switzerland. Minimum annual values extracted from that database were fitted to an extreme value distribution (Gumbel distribution) with the R package extRemes (Gilleland & Katz, 2016) to obtain the return period. *Heat wave*: +5-11 °C maximum temperature for a week, temperature range that matches the predictions of increased maximum annual temperature for Switzerland (CH2018, 2018; MeteoSwiss & ETH Zurich, 2025). To apply the heat wave, selected mesocosms were transferred to a nearby greenhouse without temperature control for the week. Therefore, temperature depended on weather conditions, and it differed between batch 1 and batch 2 (Supplementary Fig. 2). Batch 1 received +5.0 °C daily maximum temperature in average, batch 2 +11.1 °C. We accounted for this difference between the batches during our statistical analyses.

All fertilisers and chemicals were applied as liquids in a total volume of 50 ml per pot. All were dissolved in water, except for the insecticide and the fungicide. For the insecticide, 6.8 mg of thiamethoxan were diluted in 3 ml of acetone and then mixed with 80 ml of water. In case of the fungicide, 10.6 mg of carbendazim was dissolved in 2 ml ethanol and mixed with 40 ml water. Both solutions were left open overnight to allow for the evaporation of the organic solvent. Solutions were mixed per treatment combination and diluted up to 50 ml with water before application to allow the chemicals to spread throughout the soil. N and P doses were divided into two and applied at the beginning of the treatments and two weeks after. Pesticides and antibiotic were applied only once. At the time of the application, all mesocosms not receiving any chemicals received 50 ml of water.

Perturbation treatments were not applied in a full-factorial way but instead we used different levels of “perturbation richness”, namely 2, 5, 8 and 10 perturbations, using random sampling without replacement from the pool of 10 perturbations. Therefore, individual combinations were not replicated, but each replicate was a different combination of perturbations. We created ten replicates for each perturbation richness level. All the perturbations were also applied individually (1 perturbation), with eight replicates each. 20 mesocosms were kept as control and did not receive any perturbations. In total we had 140 mesocosms, 70 in each batch. This design, while maintaining the number of samples manageable in terms of logistics and manpower, allowed us to assess if and how the number of perturbations impacts ecosystem responses. Moreover, having all perturbations applied individually allowed us to model the response of the combined perturbations based on different null models and assess if the observed effects of multiple perturbations on ecosystem functions and properties are additive, dominative, synergistic or antagonistic (Rillig et al., 2019). We acknowledge that some perturbations are more likely to occur together than others, e.g., defoliation with trampling, or heat wave with drought. However, as all perturbations used in this study are common management and within expected climatic change scenarios, all the combinations used could realistically occur under field conditions.

### Pre-treatment soil measurements

At the first mesocosm collection date, we also collected soil samples to characterise physical and chemical soil properties (Table Supplementary 2). Coarse fraction bulk density was assessed by collecting five undisturbed soil cores (5 cm Ø, 15 cm depth), drying the material at 105 °C for 48 h and weighing it. An additional five soil cores of the same dimension were collected, dried at 60 °C and sieved through 2 mm mesh. Soil texture was determined using the sedimentation method according to Gee and Bauder (1986). Soil pH was measured potentiometrically in a suspension of 0.01 M CaCl_2_ (Soil:solution = 1:2; Electrode: Bioblock; pH meter 691, Metrohm, Switzerland). A subsample of soil was ground into fine powder (< 50 µm) in a ball mill (MM 400, Retsch, Germany) to determine total C and N content with an elemental analyser (NC 2500, Carlo Erba Instruments, Germany), and soil organic matter (SOM) content by loss on ignition (4 h at 550 °C) (Sparks, 1996).

### Post-treatment measurements

#### Ecosystem response variables

Ecosystem gas exchange of each mesocosm was measured with an infra-red gas analyser (IRGA; EGM-4, PP systems, USA; closed system) attached to a custom-made plexiglass chamber that enclosed the entire mesocosm (including the plants; 40 cm high x 30 cm diameter cylindrical chamber). The chamber had a total volume of 23.77 L excluding the mesocosm and had a fan installed inside for air mixing. After 3 min stabilisation time upon closing the system, changes in CO_2_ concentration within the chamber were recorded every 15 s for 2 min. Measurements were conducted under ambient light (average photosynthetic active radiation (PAR) was 410 µm s^-1^ m^-2^) and full dark conditions one day before we harvested the mesocosm. We calculated net CO_2_ fluxes based on both respiration and photosynthesis under full light conditions [hereafter net CO_2_ flux (mg CO_2_ m^-2^ h^-1^)], and ecosystem respiration [plant and soil, hereafter respiration (mg CO_2_ m^-2^ h^-1^)] based on measurements taken in full dark. The CO_2_ change rate (ΔCO_2_) was calculated fitting linear models to the data of each mesocosm and extracting the slope. We then calculated the CO_2_ flux (Dossa et al., 2015) using: CO_2_ flux (µmol m^-2^ s^-1^) = ΔCO_2_ * (P*V) / (R*T*S), P being the atmospheric pressure (KPa), V the air volume inside the chamber (m^3^), R the universal gas constant, T the air temperature inside the chamber (K) and S the surface of the mesocosm (m^2^). This value was further transformed into mg CO_2_ m^-2^ h^-1^. Temperature and atmospheric pressure inside the CO_2_ chamber were continuously recorded with a Kestrel D3 Data Logger (Kestrel Instruments, PA, USA).

Organic matter (OM) decomposition rate was evaluated using the tea bag index (Keuskamp et al., 2013), with modifications to adapt to the shorter than standard time of our experiment (seven instead of twelve weeks). One green teabag (Lipton EAN: 8722700055525) and one rooibos teabag (Lipton EAN: 8711327514348) (pre-weighted) were buried in each mesocosm at around 4 cm depth twelve days after mesocosm collection. At harvest (total of 48 days of incubation period), the teabags were carefully recovered and dried at 60 °C for 48 h. Afterwards, bags were cut open and the remaining tea was weighed. Bags that were broken and had lost material were discarded. Using the equations from Keuskamp et al., (2013), we calculated *S* (stabilisation factor) and *k* (decomposition rate), as follows:

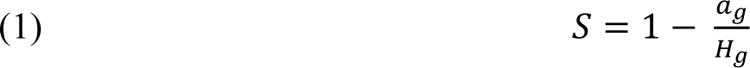

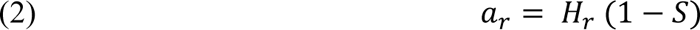

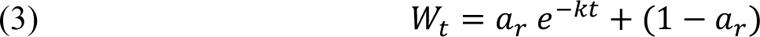

*a*_g_ is the decomposed fraction of green tea (1 – final weight green tea / initial weight green tea), *H*_g_ the hydrolysable fraction of green tea set as 0.842, *a*_r_ the decomposable fraction of rooibos tea, *H*_r_ the hydrolysable fraction of rooibos tea set as 0.552, and *W*_t_ the remaining fraction weight of rooibos tea after time *t*.

Aboveground plant biomass was cut at harvest and separated into green and brown (dead) material. Dry weight was obtained after drying at 60 °C for 48 h. Plant biomass cut as part of the defoliation treatment was also dried and weighed and added to the total aboveground biomass of the defoliated mesocosms.

#### Potential soil enzymatic activities

The potential activity of eight soil enzymes involved in OM decomposition and C, N and P cycling was measured. C-related enzymes were *β*-glucosidase (GLC) and cellobiohydrolase (CBH) that degrade cellulose, xylosidase (XYL) that degrades hemicellulose, phenoloxidase (POX) and peroxidase (PER) that oxidise lignin. *N*-acetylglucosaminidase (NAG) degrades chitin and therefore it is involved in both C and N cycling. Leucine aminopeptidase (LAP) is involved in peptide degradation (N cycling). Finally, acid phosphatase (PHO) which degrades organo-phosphate complexes, was quantified as a measure to assess P cycling.

GLC, CBH, XYL, NAG, LAP, and PHO were measured fluorometrically, according to Saiya-Cork et al. (2002), with modifications. 2 g of fresh sieved (4mm) soil stored at 4 °C and processed within 4 days of harvest were suspended in 60 ml of Tris buffer (100 mM, pH 7.0). 200 μl of soil slurry were extracted under continuous shaking, pipetted into a 96-well black plates and mixed with 50 μl of a saturating substrate-analogue solution: 1000 µM 4-MUB-*β*-D-glucopyranoside for GLC, 500 µM 4-MUB-*β*-D-cellobioside for CBH, 750 µM 4-MUB-*β*-D-xylopyranoside for XYL, 1500 µM 4-MUB-N-acetyl-*β*-D-glucosaminide for NAG, 1500 µM L-Leucine-7-amino-4-methylcoumarin for LAP, and 2000 µM 4-MUB-phosphate for PHO, all dissolved in a 2% methylcellosolve solution. Sample, substrate and quenching controls were added. Plates were incubated at 25 °C for 2 h in the dark, under continuous shaking. After the incubation period, 10 µl of NaOH 0.5 N were added to each well to increase the pH and stop the reaction. Exactly 1 min after NaOH addition, fluorescence was measured (365 nm excitation wavelength, 445/450 emission wavelength for MUB and AMC respectively) in a TECAN Infinite M200 Plate reader (TECAN, Switzerland). Reported enzyme activities are the mean of four analytical replicates.

Oxidases activity (POX and PER) was measured photometrically according to Sinsabaugh & Linkins (1988) with modifications. 0.2 ml of the same soil slurry as above were extracted under continuous shaking and mixed (1:1) with a 20 mM _L_-3,4-dihydroxyphenylalanin (_L_-DOPA) solution in a deep-well block. Blocks were shaken for 10 min and centrifuged (1000 *g*, 3 min). 200 µl of the supernatant were pipetted into transparent 96-well plates. For peroxidase activity, wells additionally received 10 µl of a 0.3% H_2_O_2_ solution. Absorbance was measured at 460 nm in the plate reader (TECAN Infinite M200) at this point (starting point) and after incubation at 25 °C for 2 h. Enzyme activity was calculated from the difference in absorption between the two time-points divided by L-DOPA molar extinction coefficient (7.9 µmol^-1^ DeForest, 2009), the path length, the amount of soil incubated in each well and the time. Reported enzyme activities are the mean of four analytical replicates.

All potential enzyme activities were expressed in nmol of product per hour per gram of dry soil (nmol MUB/AMC/oxidised DOPA h^-1^ g^-1^).

#### Soil nutrients and microbial biomass

Dissolved organic carbon (DOC) and inorganic carbon (IC) were assessed from water extracts (5 g fresh soil in 35 ml MiliQ water). Soil with extracting solution was shaken in an end-over-end shaker for 10 min, centrifuged for 30 min at 5000 rpm, and filtered through a 0.45 µm syringe filters. C was measured in a TOC-L analyser (Shimadzu, Japan). Plant available N (ammonium and nitrate) was measured based on KCl extracts (5 g fresh soil in 25 ml 1M KCl). Samples were shaken for 1 h as above and filtered through ashless filter paper (DF 5895 150, Hahnemühle FineArt GmbH, Germany). NH_4+_ was assessed colorimetrically by Berthelot reaction (Krom, 1980), and NO_3_ ^-^ by Griess reaction after reduction to nitrite with VCl_3_ (Doane & Horwáth, 2003). Soil water and KCl extracts were prepared within 4 days after harvest using fresh soil sieved through a 4 mm sieve and stored at 4 °C.

Phosphate was extracted with sodium bicarbonate solution (0.5 g dry soil + 30 ml 0.5 M NaHCO_3_). Samples were shaken overnight (16 h), then centrifuged for 10 min at maximum speed. Supernatant was pipetted into new tubes. Phosphate concentration was evaluated colorimetrically with the malachite green method (Rahutomo et al., 2019).

Microbial biomass C and N were measured using the fumigation–extraction technique (Brookes et al., 1985; Vance et al., 1987) from fresh 4 mm sieved soils, within two weeks after harvest. 5 g of soil were fumigated with CHCl_3_ for 24 h. Soluble C and N were extracted from the fumigated and un-fumigated samples with 25 ml 0.5 M K_2_SO_4_. The soil and K_2_SO_4_ were shaken for 30 min and filtered (DF 5895 150, Hahnemühle FineArt GmbH, Germany). Total C and N were analysed in diluted (1:10 in H_2_O) and acidified (with 3M HCl) samples in a TOC-V (Shimadzu, Japan). Microbial C and N flush (difference between fumigated and un-fumigated samples) were converted to microbial biomass using k_EC_ factor of 0.35 (Sparling et al., 1990) and k_EN_ factor of 0.54 (Brookes et al., 1985).

#### Soil physical properties

Water surface infiltration time was used as a proxy for soil water repellency (Hallett, 2007). To do that, we placed a drop of de-ionised water (8 μl) on the soil surface and recorded the time (in seconds) that it needed to completely infiltrate into the soil. Time was recorded up to 2 minutes, and if drops were not infiltrated by then, > 120 s was recorded instead. Three independent measurements were done per mesocosm, which were averaged for statistical analyses (technical replicates).

Water stable aggregates were evaluated according to Kemper and Rosenau (1986). Briefly, the percentage of water stable aggregates was determined by placing 4 g of dry soil (dried at 40 °C), on 0.25 mm mesh size sieves. After capillary re-wetting with deionized water, samples were inserted into a sieving machine (Agrisearch Equipment, Eijkelkamp, Giesbeek, Netherlands) and sieved for 3 min. Percentage of water-stable aggregates (%WSA) per sample were calculated as: %WSA = (water stable fraction-coarse matter)/(4.0 g-coarse matter).

### Data analyses

Initially, all variables were analysed with linear models including “Treatment” (with 15 levels, all the perturbations applied individually plus the combinations of 2, 4, 6, 8 and 10 simultaneous perturbations) and “Batch” as fixed factors. For enzyme data, “Enzyme plate” was used instead of “Batch”. This is because “Enzyme plate” is nested in “Batch” (plates 1-3 belonged to batch 1 and 4-6 to batch 2) and “Enzyme plate” allows us to better control for the small changes due to measurement date. Box-Cox transformations were used to achieve normality of the residuals when needed, as detailed in Supplementary Table 3. To account for different variance in heteroskedastic variables, we added the term “weights” to the models using the inverse of the variance per group where needed.

Then, different null models were used to predict the response of our variables to multiple combinations of perturbations based on their response to individual perturbations (Fig. 1). The additive null model was calculated as the addition of the estimated coefficients of the different individual perturbations with a linear model, including the batch (or the enzyme plate for enzyme variables) as fixed factors, and the term “weights” as above. To avoid predictions becoming negative, a negative binomial model was used instead for respiration and water surface infiltration time. The function *add_ci* from the package “ciTools” (version 0.6.1; Haman and Avery, 2020) was used afterwards to predict the 95% confidence intervals of the predictions of the null models. Any values higher than the upper confidence interval (95%) of the additive model were considered as synergistic responses. Any values lower than the lower confidence interval (5%) were considered as antagonistic responses. For visualisation, the predictions of the models and their confidence intervals were scaled between −1 and 1, 0 being the prediction of the model. To account for the different directions of effects sizes, we modified the sign of the rescaled value so that if the additive model predicts a decline, a further decline represents a synergy (positive sign in the rescaled value), while a smaller decline represents an antagonistic response (negative sign).

Besides the additive model we also calculated dominative and proportional models. For the dominative model we used the effect of the highest absolute value of the individual coefficients (Fig. 1). This model will fall within the antagonistic zone when all factors have the same direction but can fall within the synergistic zone if the individual factors show opposite effects (Fig. 1C). The proportional model was calculated by dividing the estimated coefficients by the number of perturbations applied simultaneously (Fig. 1). This model will always fall within the antagonistic zone. Confidence intervals for these models were calculated as above for the additive model. Finally, all variables were combined in a principal component analysis (PCA) using the *rda* function from vegan (version 2.6-4; Oksanen et al., 2024) to assess the functional response of the soils to the different treatments. Finally, to disentangle the effect of the presence/absence of each perturbation compared to the effect of the number of perturbations in the measured response variables, we carried out ten independent linear models with the perturbation to be tested as a binary yes/no variable, the number of perturbations, their interaction, and batch or enzyme plate as fixed factors, including the term “weights” as above, for each response variable. This analysis helps to identify which perturbations have stronger dominative effects. All analyses were done in R v 4.4.1 (R Core Team, 2025).

## Results

Most of the soil variables, plant variables and ecosystem response variables measured were strongly affected by the individual perturbations and the different combination of multiple perturbations (Fig. 2, Fig. S3), showing in some cases additive effects or dominative effect of the drought treatment (Fig. 3, 4). However, against our expectations, we did not detect clear synergistic effects.

**Fig. 2:**
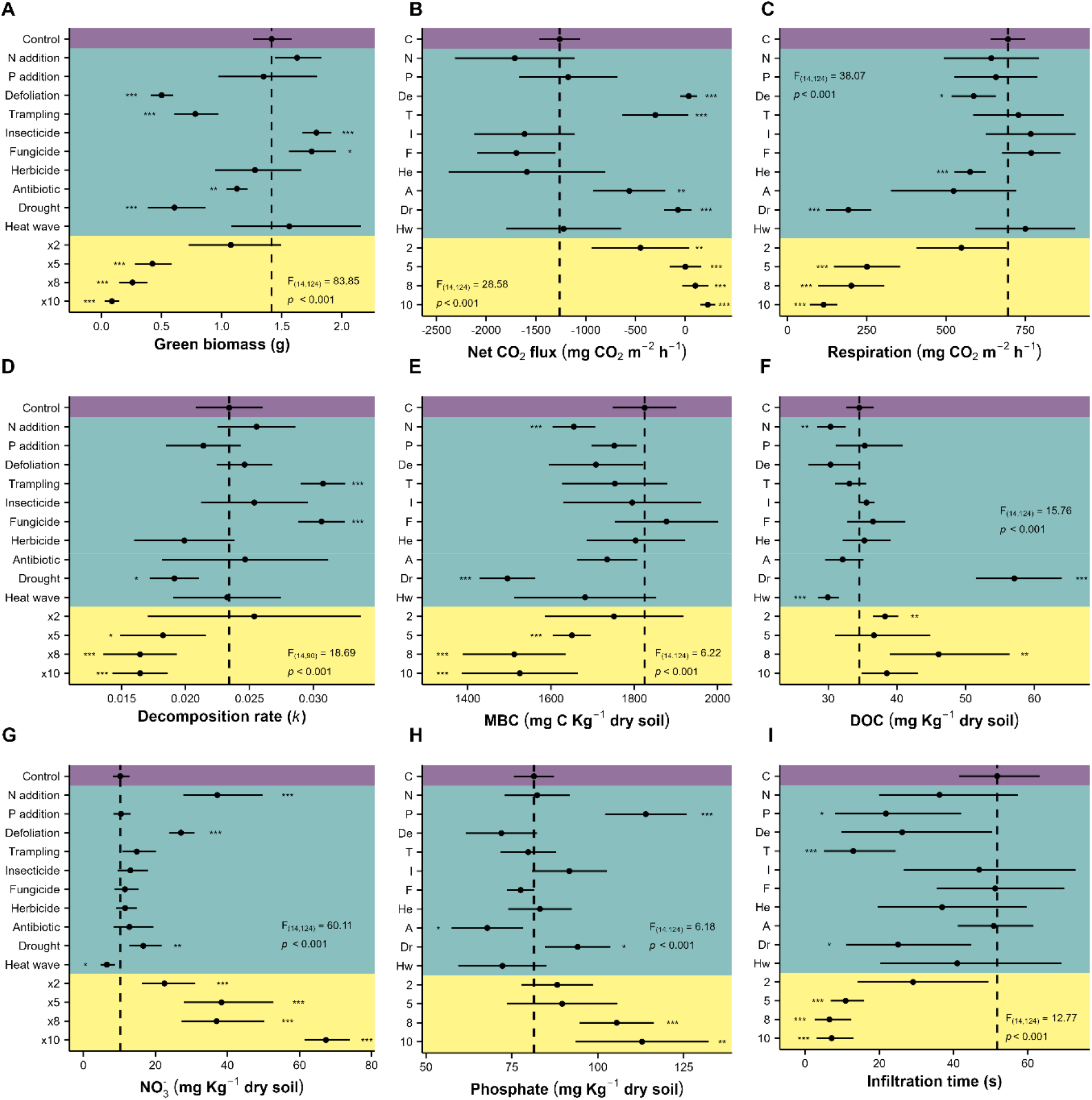
Effects of individual and combined perturbations on the different response variables. MBC = microbial biomass carbon, DOC = Dissolved organic carbon. Estimated marginal means of the model plus 95% interval confidence are shown. Significance of the effect is symbolised as °: p < 0.1, *: p < 0.05, **: p < 0.01, ***: p < 0.001.

**Fig. 3:**
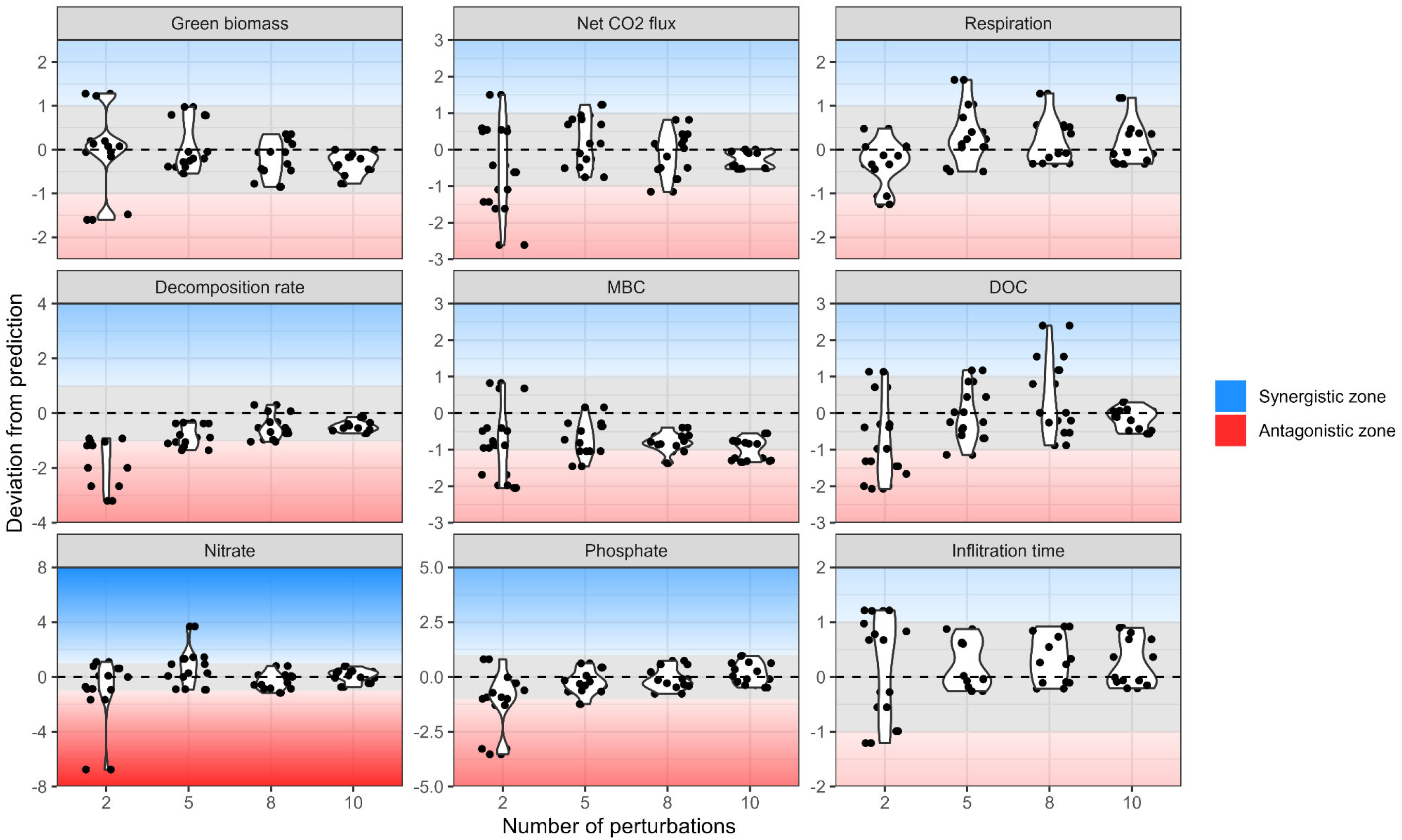
Deviation from the prediction of the additive model of combined perturbations for each response variable. The grey zone indicates the 95% confidence interval of the on. Points that fall above the grey area are in the synergistic zone and points that fall below the grey area are in the antagonistic zone. For variable units see Fig. 2. MBC = ial biomass carbon, DOC = Dissolved organic carbon.

Plant green biomass was reduced by defoliation (−65%), trampling (−45%), antibiotics application (−20%), and drought (−57%), while it increased after insecticide (+26%) and fungicide applications (+24%) (Fig. 2A). When more than two perturbations were applied simultaneously, plant green biomass was reduced in all cases (−23%, −70%, −82%, and −94% for 2,5, 8 and 10 combined perturbations, respectively, Fig 2A). In contrast, plant brown biomass was only significantly affected by P addition (−24%, Fig. S3A). Total plant biomass (green and brown) was positively affected by N addition (+18%) and negatively by trampling (−21%), herbicide addition (−15%) and combinations of at least five perturbations (−25%, −24%, and −18% for 5, 8 and 10 combined perturbations, respectively; Fig. S3B).

Net CO_2_ flux became more positive (i.e., higher C losses) with defoliation (+103%, changed from C assimilation to emission), trampling (+76%), antibiotic addition (+55%), drought (+94%) and in all the combinations of perturbations (+64%, +100%, +108%, +118%, for 2, 5, 8 and 10 combined perturbations, respectively) (Fig. 2B). In parallel, ecosystem respiration declined with defoliation (−16%), herbicide application (−17%), and specially with drought (−72%) and all the multiple perturbations (except for the combinations of 2 perturbations, −64%, −71%, and −84%, for 5, 8 and 10 combined perturbations, respectively) (Fig. 2C).

Decomposition rate (*k*) was enhanced by trampling and fungicide application (+31% each) and declined under drought (−18%) and the combination of five (−22%) or more perturbations (−30% for both 8 and 10 combined perturbations; Fig. 2D). The stabilisation rate (*S*) was significantly reduced by N addition (−12%) and antibiotic application (−15%), and increased under drought (+47%) and combinations of five or more perturbations (+7%, +34%, and +28% for 5, 8 and 10 combined perturbations; Fig. S3C).

Microbial biomass C (MBC) showed a significant reduction after N addition (−9%), drought (−18%), and combinations of five or more perturbations (−10%, −17%, and −16% for 5, 8 and 10 combined perturbations; Fig. 2E). Microbial biomass N (MBN) was reduced by the same factors plus by antibiotic addition (−9 % N addition, −44% drought, −6% antibiotic addition, −31% 5 perturbations, −36% 8 perturbations, −30% 10 perturbations; Fig. S3D). The microbial C:N ratio was significantly reduced by defoliation (−4%) while it strongly increased with drought (+40%) and combinations of at least 5 perturbations (+16%, +28%, and +20% for 5, 8 and 10 combined perturbations; Fig. S3E).

DOC significantly decreased after N addition (−12%) and heat wave (−13%), while it showed a strong increase with drought (+65%; Fig. 2F). Combinations of two or eight perturbations increased DOC values (+11%, +33%, respectively). IC was significantly reduced by all the combinations of perturbations (−11%, −19%, −21%, and −28% for 2, 5, 8, and 10 combined perturbations) but not for any of the individual perturbations (Fig. S3F). Plant available nitrate was reduced by the heat wave (−36%), and increased after N addition (263%), defoliation (+164%), drought (+62%) and all the combinations of perturbations (+120%, +274%, +261%, and +558% for 2, 5, 8 and 10 combined perturbations; Fig. 2G). Plant available ammonium was not strongly affected by the experimental treatments but significantly decreased after P addition (−6%) and herbicide application (−8%) and increased after insecticide addition (+10%) and when applying all ten perturbations simultaneously (+9%) (Fig. S3G). Available phosphate increased with P addition (+40%), drought (+16%), and combinations of eight (+30%) or ten (+39%) perturbations, while it decreased with antibiotic addition (−17%) (Fig. 2H). Soil pH only slightly varied with experimental treatments, increased significantly with antibiotic addition (+1%) and declined with five perturbations applied simultaneously (−1%) (Fig. S3H).

Soil enzymatic activities were not strongly affected by experimental treatments (Fig. S4). Only a significant effect of herbicide addition was observed for the activity of CBH (+20%) and NAG (+25%) enzymes (Fig. S4). Finally, water surface infiltration time decreased with the different perturbations, significantly after P addition (−58%), trampling (−75%), drought (−52%), and combinations of five or more perturbations (−79%, −87%, and −86% for 5, 8 and 10 combined perturbations; Fig. 2I). Percentage of water stable aggregates was not affected by any of the experimental treatments (Fig. S3I).

The combined effect of multiple perturbations could be partially predicted based on the effects of individual perturbations, as the observed values fell within the limits of the additive model (Fig. 3). However, the predictions had large confidence intervals which became larger when the number of simultaneous perturbations increased (Fig. S5), making the predictions more uncertain as we increased the number of simultaneous perturbations. Due to these large confidence intervals, the observed values could in many cases also be explained by the dominative or the proportional model (Fig. S6).

When only two perturbations were applied, we observed in many instances antagonistic effects (Fig. 3). These antagonistic effects did, however, disappear as we increased the number of perturbations, probably due to the increased uncertainty of our models (Fig. S5). However, microbial biomass variables (MBC and MBN) consistently showed antagonistic effects, and the data were better explained by the proportional model (Fig. 3, Fig. S6, Fig S7). On the other hand, respiration and DOC responses showed some synergistic effects, but not consistently (Fig. 3). Although still within the limits of the additive model, green biomass and net CO_2_ flux were better explained by the dominative model (closer to 0 in Fig. S6). Plant available nitrate showed a lot of variation with many points falling outside the additive zone when 2 or 5 perturbations were applied, response which was not explained by any of the explored models. However, it was better explained by the additive model than the dominative or the proportional models when 8 or more perturbations where applied (Fig. S6). The rest of the variables were equally explained by any of the three tested models (Fig. S6, S7).

The number of perturbations applied significantly affected many of the measured response variables (Supplementary Table 4), but on some occasions the presence of a specific perturbation had specifically strong effects. This was especially true for drought. For example, for the stabilisation factor (*S*), MBC, MBN, microbial C:N ratio, and DOC, we only found a significant main effect of the presence/absence of the drought, but no significant effect of the number of perturbations. These findings therefore strongly suggest a dominative effect of drought (Table Supplementary 4), which was also underpinned by our ordination analysis (PCA; Fig 4). The mesocosms clearly separated according to the presence/absence of drought (Fig. 4). This pattern was not observed when points were plotted by shape by other perturbations, just with drought.

**Fig. 4:**
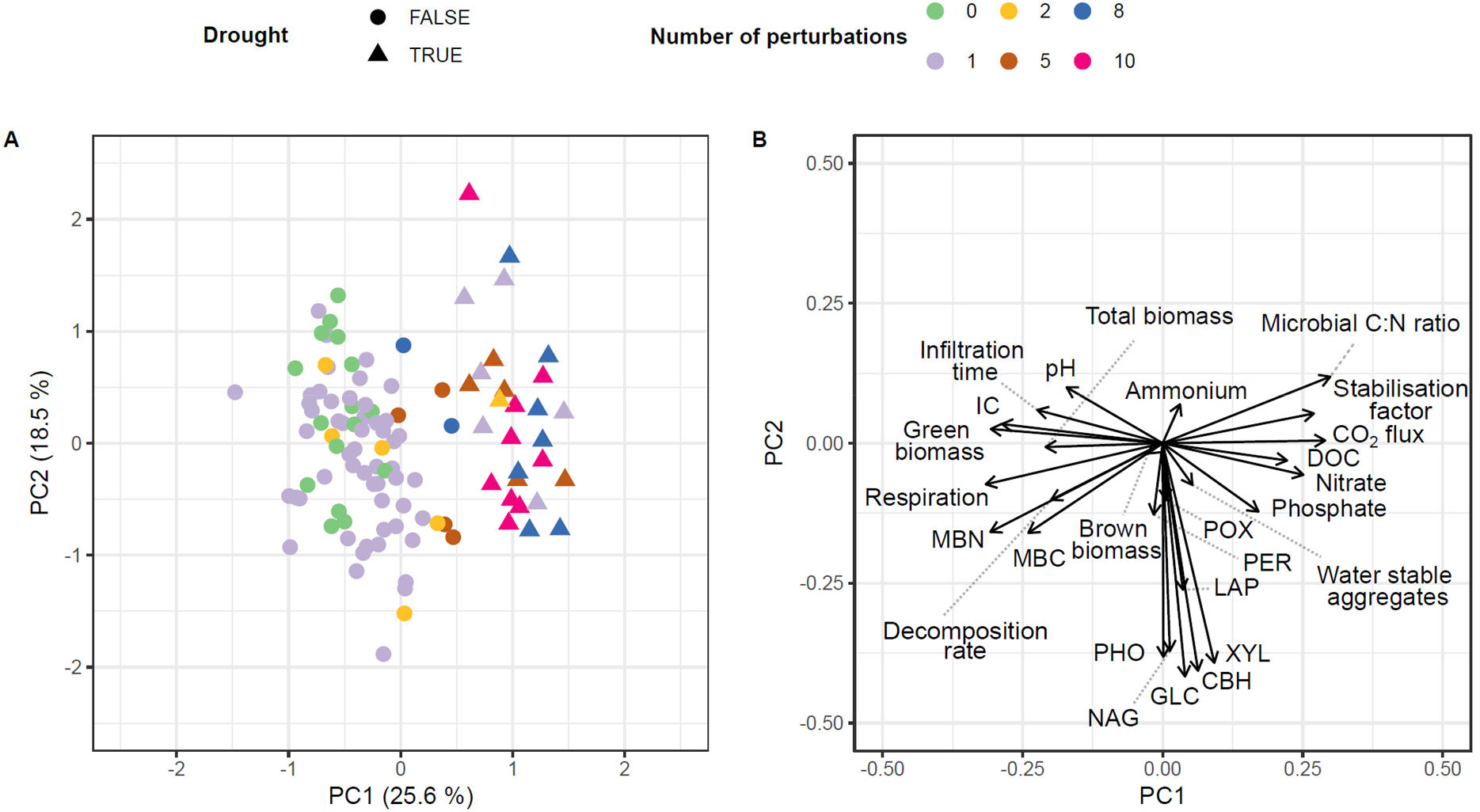
**a)** Principal component analysis (PCA) including all the measured response variables. b) Variable loading vectors for all response variables.

## Discussion

Individual perturbations had strong and diverse effects on grassland functionality. However, directional negative effects were present when increasing number of perturbations were applied simultaneously. For many of our variables measured, we observed additive effects of the combined perturbations. Nevertheless, drought often resulted in dominative effects masking the effect of other perturbations. Further, we detected some antagonistic effects of the different perturbations. However, against our expectations, we did not detect clear synergistic effects. This rather surprising result could potentially be related to a buffering effect of the plant community, which we will discuss further below.

### Effects of individual perturbations

Drought was the perturbation with the strongest individual effect on ecosystem functioning and properties in this experiment and also drove many of the observed effects of the combined perturbations in a dominative manner. This is similar to what has been reported previously for grassland functionality (Barnard et al., 2013; Cordero et al., 2023; C. Liu et al., 2023; Ochoa-Hueso et al., 2018), e.g., negative drought effects on plant biomass (C. Liu et al., 2023), net C uptake (Thompson et al., 2020), microbial biomass (Hueso et al., 2012; Preece et al., 2019), and decomposition rates (Deng et al., 2021). Positive drought effects were in contrast found for soil nutrient availability due to microbial death and reduction of plant uptake (Deng et al., 2021). Additionally, bacteria have been shown to be more sensitive to drought than fungi (Bapiri et al., 2010; Canarini et al., 2024; de Vries et al., 2018), which could explain the observed increase in the microbial C:N ratio with drought in our study (Strickland & Rousk, 2010).

Nutrient enrichment, through N deposition or mineral fertilisation, can have strong impacts on ecosystems (Borer & Stevens, 2022; Fowler et al., 2013; Peñuelas et al., 2013). As other studies, we found that N addition increased plant biomass (Borer & Stevens, 2022; LeBauer & Treseder, 2008) and decomposition rates (Ochoa-Hueso et al., 2020), especially at early decomposition stages (Gill et al., 2022). Also similar to earlier findings we found reduced microbial biomass after fertilisation(Farrer et al., 2013; Treseder, 2008). However, we also observed a decline in DOC, against general trends of increased DOC with fertilisation (C. Xu et al., 2021). P additions reduced brown plant biomass, which suggests that the plants were healthier in mesocosms that received P, which potentially released P limitation for plant growth (Vitousek et al., 2010).

Defoliation of plants is known to exert strong effects on soil properties (Dollete et al., 2024; N. Liu et al., 2015; Y. Liu et al., 2023) and ecosystem respiration (Iqbal et al., 2012; Skinner & Goslee, 2016). The later are highly variable as ecosystem respiration is related to both plant and soil microbial respiration. Defoliation was often linked to an increased microbial growth as defoliated plants increase root exudation (Hamilton et al., 2008; T. Xu et al., 2024). However, in our study, we could not observe a significant effect of defoliation on microbial biomass, as also found for grasslands globally (Risch et al., 2023), but we did observe a reduction in the microbial C:N ratio. This reduction could be linked to i) a shift towards a bacterial dominated microbial community, as bacteria can quickly benefit from the labile C produced when root exudations increase (Shahzad et al., 2012; Williamson & Wardle, 2007), or ii) a C limitation compared to N as described in other studies (Medina-Roldán & Bardgett, 2011). It is known that soil available N increases after defoliation (Capstaff et al., 2021; Dollete et al., 2024; Y. Liu et al., 2023) as defoliation stimulates N mineralisation (Medina-Roldán & Bardgett, 2011), and reduces plant uptake (Shahzad et al., 2012).

Trampling by cattle can have highly negative effects for soils in pastures (Centeri, 2022; Trimble & Mendel, 1995), for example, by decreasing plant biomass as observed in our experiment (Dunne et al., 2011; Hiltbrunner et al., 2012), that resulted in lower C assimilation of our trampled mesocosms. The increase in the decomposition rate *k* observed could be explained by the fact that trampling tends to affect the soil invertebrate community more negatively than the soil microbial community (Larsen et al., 2004; Lee et al., 2009). As invertebrates, such as collembola, feed on microorganisms (Potapov et al., 2022), a reduction in their abundance may therefore lead to a release of feeding pressure on microbial communities and therefore to an increase in decomposition rates. Finally, we observed a reduction in the water surface infiltration time, i.e., faster infiltration rates in trampled soils. This is against expectations as soil compaction usually reduces infiltration rates (Nawaz et al., 2013). However, this effect could be related to changes in the biofilm layer in the soil surface. Control undisturbed mesocosm likely had a biofilm on the soil surface, that created soil water repellency (Epstein et al., 2011; Schaumann et al., 2007). Trampling could have physically destroyed this layer, increasing water surface infiltration (Kuske et al., 2012).

Pesticide applications had only mild effects on our measured variables. They increased plant green biomass (insecticide and fungicide), as expected, as they released the plant community from pathogens (Bai et al., 2024; Simon-Delso et al., 2015). Also as expected, herbicide addition reduced total aboveground biomass and therefore reduced ecosystem respiration. Additionally, herbicide application increased the activity of CBH and NAG enzymes, which are known to be involved in the degradation of OM by microbial communities (Ai et al., 2023). Further, fungicide increased the decomposition rate *k*, which could point toward a change in competition between bacteria and fungi (Hicks et al., 2019; Mille-Lindblom & Tranvik, 2003). Fungicide decreases fungal activity and growth, thus providing bacteria with better access to resources, which in turn resulted in a faster decomposition of more labile OM (Wang & Kuzyakov, 2024). Antibiotics application reduced microbial biomass, as expected (J. Chen et al., 2023), and reduced plant biomass and increased net CO_2_ flux (i.e., reduced C assimilation), either due to direct effects of antibiotics on the plants or due to the loss of beneficial effects of microorganisms for plant growth (Gworek et al., 2021).

Heat waves are known to have long-lasting effects on ecosystem functions and properties (Ummenhofer & Meehl, 2017). In our experiment, however, we only observed a significant decrease in soil nutrients (DOC and nitrate) and, although not significant, a trend towards lower activities of soil microorganisms (enzyme activities). This latter finding is supported by the fact that extreme high temperatures reduce enzymatic capacities of soils (Tabatabai, 1994), which can further lead to the reduced nutrient availability observed in our soils.

### Effects of combined perturbations

We observed consistent negative responses of our measured variables when multiple perturbations occurred simultaneously. Interestingly, this was also the case when individual responses to the perturbations differed in signs (e.g., green biomass and decomposition rate). In consequence, plant, soil and ecosystem properties had a consistently lower functionality under multiple combined perturbations. At the same time, available nutrients accumulated. These superimposed effects of multiple perturbations have been previously observed in other studies (Rillig et al., 2019). However, contrary to our expectations (Bi et al., 2024; Meidl et al., 2024; Rillig et al., 2019) in our study, we did not detect synergistic effects. We did instead observe additive, dominative and antagonistic effects. Additive effects have been commonly found in experiments with four or less perturbations applied simultaneously (Gutknecht et al., 2010; Niboyet et al., 2011) or in meta-analyses (Yue et al., 2017; Zhou et al., 2016). But most of the few studies conducted with 10 or more perturbations applied simultaneously (Bi et al., 2024; Meidl et al., 2024; Rillig et al., 2019) reported synergistic effects, which we do not find.

Antagonistic effects, i.e., effects lower than expected by the additive model, have been described in previous experiments (García-Fayos & Bochet, 2009; Henry et al., 2005; Hines et al., 2017). This antagonist response has been explained due to the fact that the single effect of one factor is so intense that when another factor is acting on the same system there is no further effect (García-Fayos & Bochet, 2009). In line with this, we observed a clear dominative effect of drought. Even a drought that is not very intense, as the one applied in this experiment (reduction to 30% WHC), can have an overpowering effect on ecosystems even in the presence of many other perturbations. This effect could have been amplified by the so-called sampling effect of drought (higher probability of including drought in the random draws with an increasing factor number of perturbations added; Speißer et al., 2022). This sampling effect is an artifact of experiments with multiple combined perturbations, but it also highlights the perturbation with strongest effects among the pool of selected perturbations. It is worth stating that the prediction intervals for all of our models widened as we increased the number of perturbations applied. This fact by itself highlights the difficulty of predicting the ecosystem responses to multiple perturbations and the importance of conducting experiments where many perturbations are applied simultaneously.

A potential explanation for the difference in results between other studies with ten or more perturbations (often synergistic effects) and ours (additive and dominative effects) could be the presence of the plants in our study, which were missing in those abovementioned experiments. Plants buffer soil and ecosystems parameters, helping the microbial communities and ecosystem functionality to recover from perturbations (Oram et al., 2025). For example, drought can modify the quantity and quality of root exudates, which then leads to the attraction of different microorganisms. Through this mechanism, interaction between plants and microbes could help the entire system to resist or recover better from drought (Williams & de Vries, 2020), and in consequence increase the ecosystem’s resilience to drought. Vegetation can also favour the persistence of richer mesofaunal communities and their top-down control on microbes, conferring additional buffering capabilities to the system (Bradford et al., 2002; Jin et al., 2022). More generally, global change perturbations can modify the direction and intensity of plant-soil feedbacks, which will ultimately regulate ecosystem functioning (Hassan et al., 2022; Pugnaire et al., 2019). We did, however, not evaluate changes in plant-soil feedbacks directly in our experiment.

Another potential reason for not observing synergistic effects in our experiment may lie in the lower bounds of the measured variables: once they approach their physiological minimum further decreases are no longer possible. Overall, our study highlights that synergistic effects are not the norm when a more holistic ecosystem approach is included in the experiment (i.e., more realistic settings), as suggested by Côté et al., (2016). We do recognize that we only conducted a five-week greenhouse experiment, and some response variables may have needed longer exposure periods to the global change perturbations applied to show a response. A meta-analysis of individual and multiple global change perturbation experiments suggested that synergistic effects are more likely to be observed in the long-term (Komatsu et al., 2019), implying that the short-term effects observed in this experiment could be amplified if the perturbations continued over time. Moreover, the fact that our different response variables responded in many different ways highlight the importance of including different response measures if we were to understand how multiple perturbations affect the ecosystem.

We acknowledge that our choice of experimental design which focussed on the replication of the number of multiple perturbations (2, 5, 8 or 10 perturbations) rather than the individual combinations of multiple perturbations (see methods) did not allow us to evaluate the effect of perturbation identity when applied in combination. Yet, our study provides a strong demonstration of the complexity and unpredictability of the effects of multiple combined perturbations on ecosystem properties and functions and paves the way for further research on the topic.

## Conclusions

Our results demonstrate that multiple combined perturbations have strong deleterious effects on grassland ecosystem functioning, effects that can only be inadequately captured by studying perturbations individually. This study also highlights the importance of studying combined perturbations under more realistic conditions than soil incubations and account for the potential buffering effect of plants. Although the studies of single individual perturbations are still crucial to gain a mechanistic understanding of the complexities of global change effects on soils, plants and ecosystem functioning, studies that combine multiple perturbations are essential to be able to predict future responses of ecosystems to continuous anthropogenic pressures. Moreover, understanding which combinations of perturbations are more detrimental for grassland functionality can guide land managers and policy makers to avoid or minimise specific management practices under climate change conditions. Future research directions should include long-term experiments, field experiments, studies on different types of grasslands, grasslands with a different background of climatic and management conditions, or experiments with different levels of the applied perturbations, among others.

## Acknowledgements

The project was funded by the British Ecological Society, grant LRB21/1006 awarded to Irene Cordero. We thank the support of the members of the Plant-Animals Interactions group, the Rhizosphere Processes group and the Soil Functions and Soil Protection group at WSL. We also thank Pedro Tognetti and Diego Abad Martín for their invaluable help collecting the mesocosms. In addition, we are thankful to the WSL garden crew for support with logistics in both mesocosm collection and the greenhouse experiment. We finally thank three anonymous reviewers for their helpful suggestions for improving the manuscript.

## Data availability

All the data are available at the Envidat database (www.envidat.ch) under https://www.doi.org/10.16904/envidat.452.

## Conflict of interest

The authors declare no competing interests.

## Author contribution

IC designed the study, with inputs from MFJ, SG, and ACR. MFJ, SG, JH, MCR, and IC carried out the experiment and lab work. ACR and SZ provided resources, personnel and expertise to carry out the project. MFJ and IC analysed the data and wrote the first draft of the manuscript. All the authors contributed to the writing of the manuscript and approved its final version. IC obtained the funding to carry out the project.

## SUPPLEMENTARY MATERIAL

**Supplementary Table 1:** List of plant species occurring in experimental soil cores and number of soil cores where it was detected, and average percentage coverage of the different plant functional groups and bare soil (estimated visually), mean ± standard deviation is shown (n = 140).

|  | Number of occurrences |
| --- | --- |
| *Ajuga reptans* L. | 29 |
| *Alopecurus pratensis* L. | 1 |
| *Anthoxanthum odoratum* L. | 3 |
| *Arrhenatherum elatius* (L.) J. & C. Presl | 56 |
| *Bromus hordeaceus* L. | 17 |
| *Centaurea jacea* L. | 1 |
| *Crepis biennis* L. | 101 |
| *Dactylis glomerata* L. | 107 |
| *Galium mollugo* aggr. | 67 |
| *Glechoma hederacea* L. | 12 |
| *Holcus lanatus* L. | 67 |
| *Lolium multiflorum* Lam. | 55 |
| *Lolium perenne* L. | 12 |
| *Lysimachia nummularia* L. | 1 |
| *Medicago lupulina* L. | 3 |
| *Plantago lanceolata* L. | 40 |
| *Poa pratensis* L. | 10 |
| *Poa trivialis* L. | 1 |
| *Potentilla reptans* L. | 49 |
| *Prunella vulgaris* L. | 5 |
| *Ranunculus acris friesianus* (Jord.) Syme | 104 |
| *Rumex acetosa* L. | 12 |
| *Tragopogon pratensis orientalis* (L.) Celak. | 8 |
| *Trifolium pratense* L. | 22 |
| *Trisetum flavescens* (L.) P. Beauv. | 33 |
| *Veronica chamaedrys* L. | 17 |
| *Veronica filiformis* Sm. | 55 |
| *Vicia sepium* L. | 61 |
|  | % coverage |
| Grasses | 59.25 ± 20.38 |
| Forbs | 41.69 ± 25.57 |
| Legumes | 4.51 ± 6.15 |
| Mosses | 27.46 ± 27.79 |
| Bare soil | 12.55 ± 11.52 |

**Supplementary Table 2:** Main field soil characteristics. Mean ± standard deviation is shown (n = 5).

| Variable | Value |
| --- | --- |
| pH | 7.01 ± 0.08 |
| Bulk density (g cm<sup>-3</sup>) | 0.92 ± 0.09 |
| Texture: |  |
| % sand | 33.7 ± 5.4 |
| % silt | 33.4 ± 2.3 |
| % clay | 32.9 ± 3.5 |
| Organic matter (%) | 15.3 ± 1.4 |
| Total C (%) | 0.54 ± 0.05 |
| Total N (%) | 6.44 ± 0.42 |
| C/N | 11.95 ± 0.54 |

**Supplementary Table 3:** Box-Cox transformations applied to the response variables for statistical analyses.

| Variable name | Transformation applied |
| --- | --- |
| Green biomass | Log +1 |
| Brown biomass | Sqrt |
| Total biomass | - |
| C flux under light | - |
| Soil respiration | - |
| Stabilisation factor (S) | 1/sqrt |
| Decomposition rate (k) | - |
| MBC | - |
| MBN | Sqrt |
| Microbial C:N | 1/x<sup>2</sup> |
| DOC | 1/x |
| IC | - |
| Ammonium | - |
| Nitrate | Log |
| Phosphate | - |
| GLC | - |
| CBH | x<sup>2</sup> |
| XYL | - |
| NAG | - |
| LAP | Log |
| PHO | - |
| POX | - |
| PER | - |
| pH | - |
| Water stable aggregates | - |
| Surface infiltration time | Sqrt |

**Supplementary Table 4:**
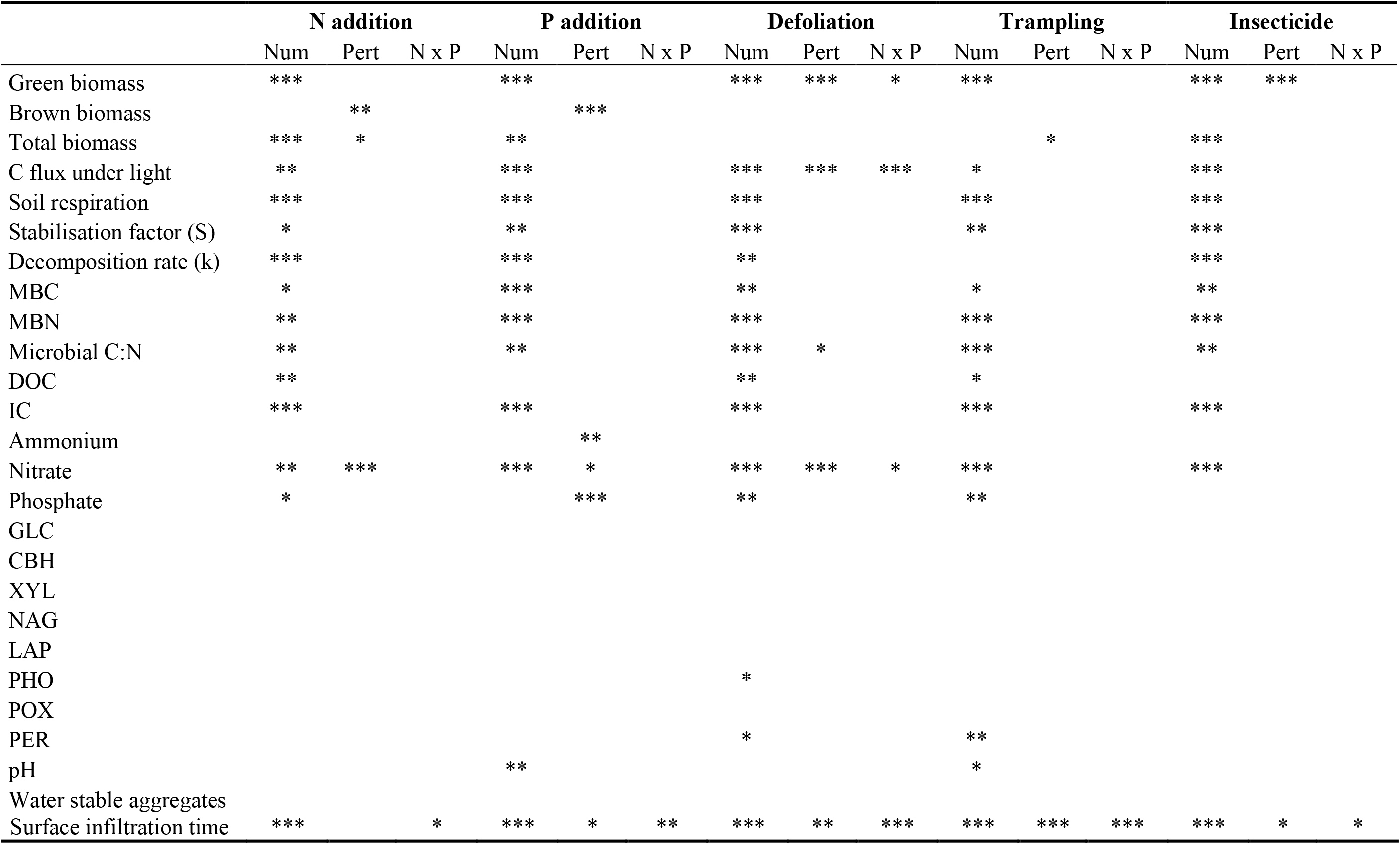
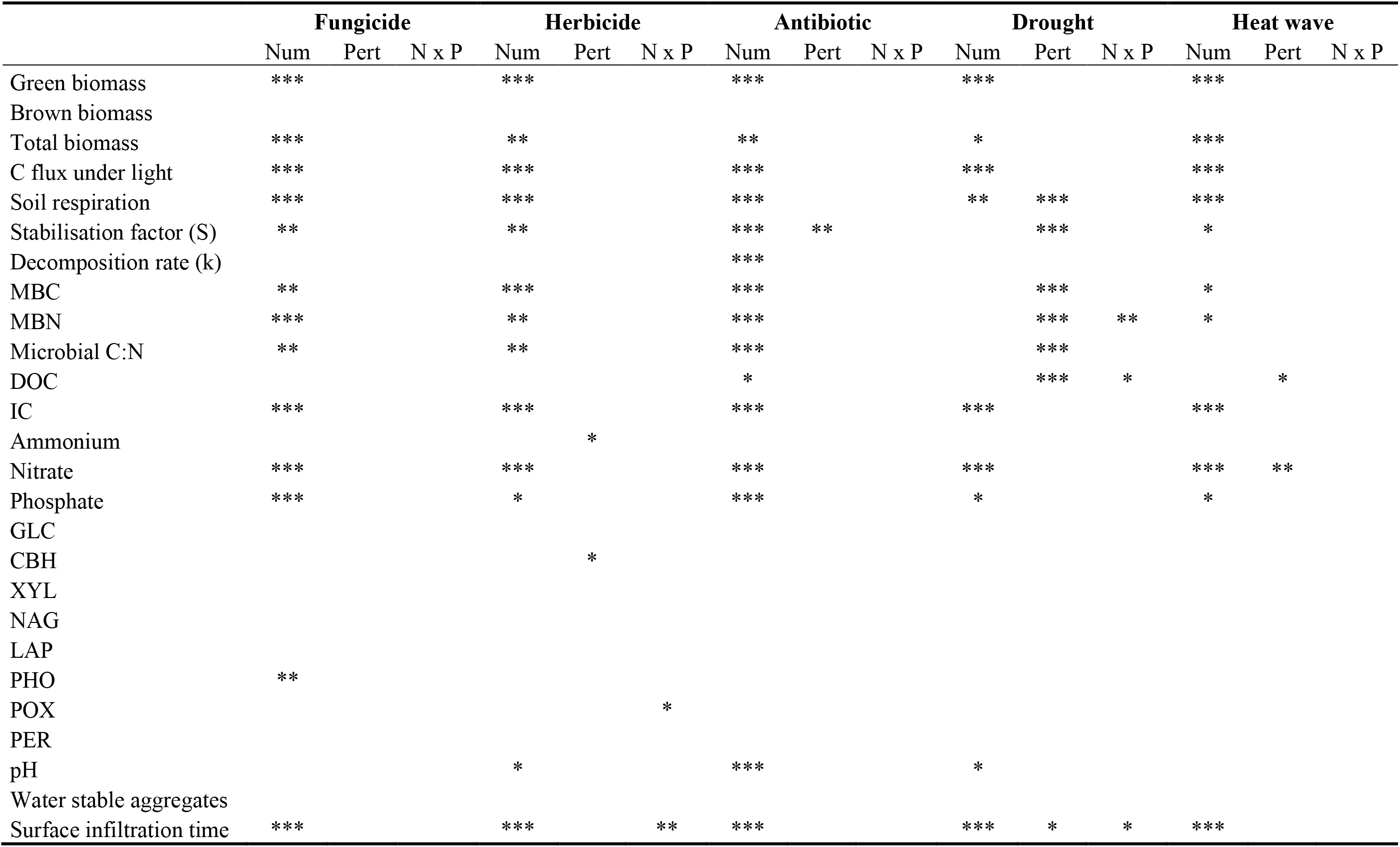
Assessment of the contribution of the presence of each perturbation to the observed effects. Linear models were calculated independently for each perturbation, where that perturbation is added as a binary yes/no variable. Benjamini-Hochberg false discovery rate adjustment was applied to correct for multiple testing. The effect of the number of perturbations (Num), the presence of the perturbation of interest (Pert) and the interaction (N x P) is shown as *: p < 0.05, **: p < 0.01, ***: p < 0.001.

---

**Fig. S1.**
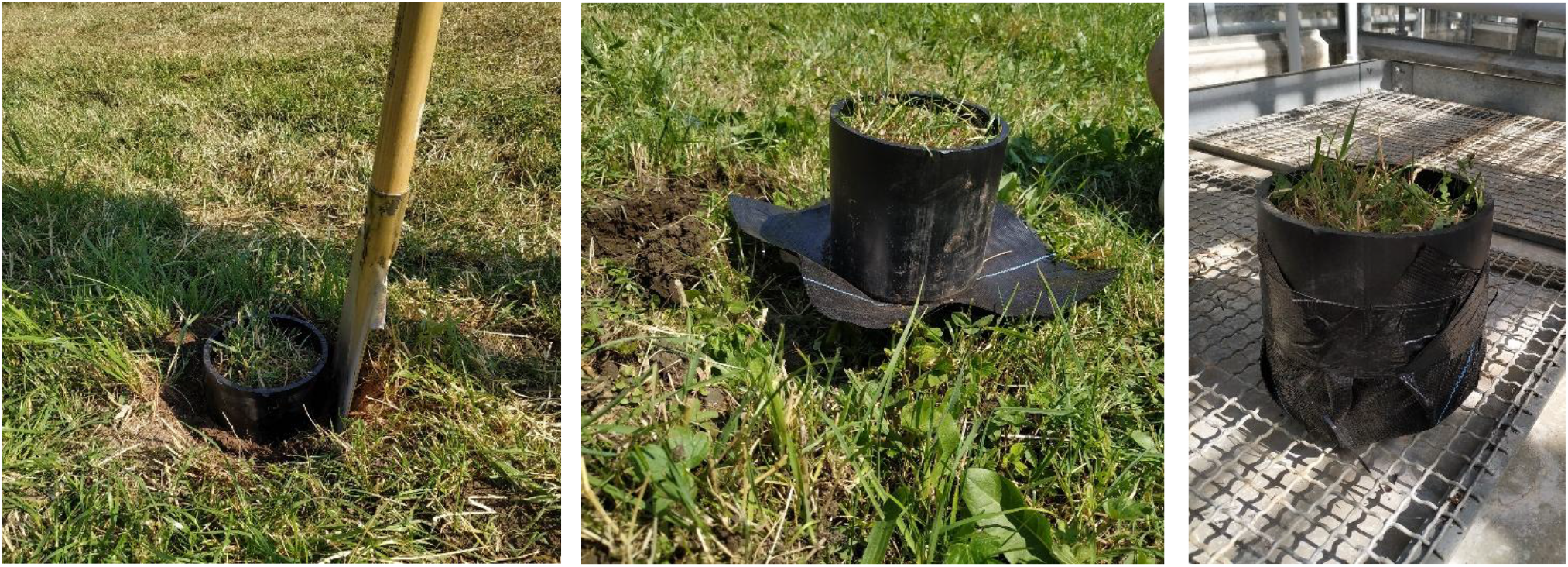
Collection of the soil mesocosms.

**Fig. S2.**
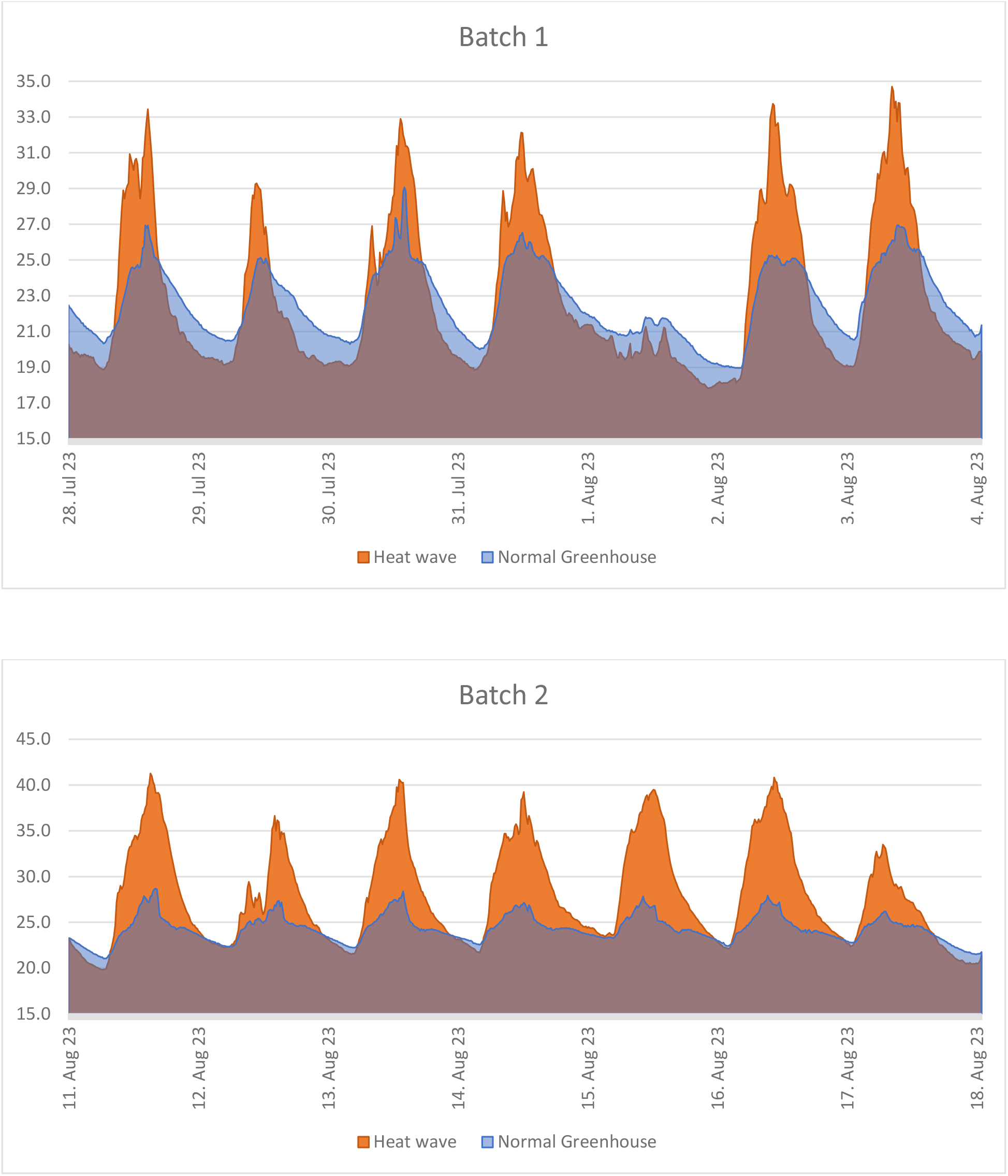
Temperature conditions in the greenhouse during the heat wave treatment in the two different batches.

**Fig. S3:**
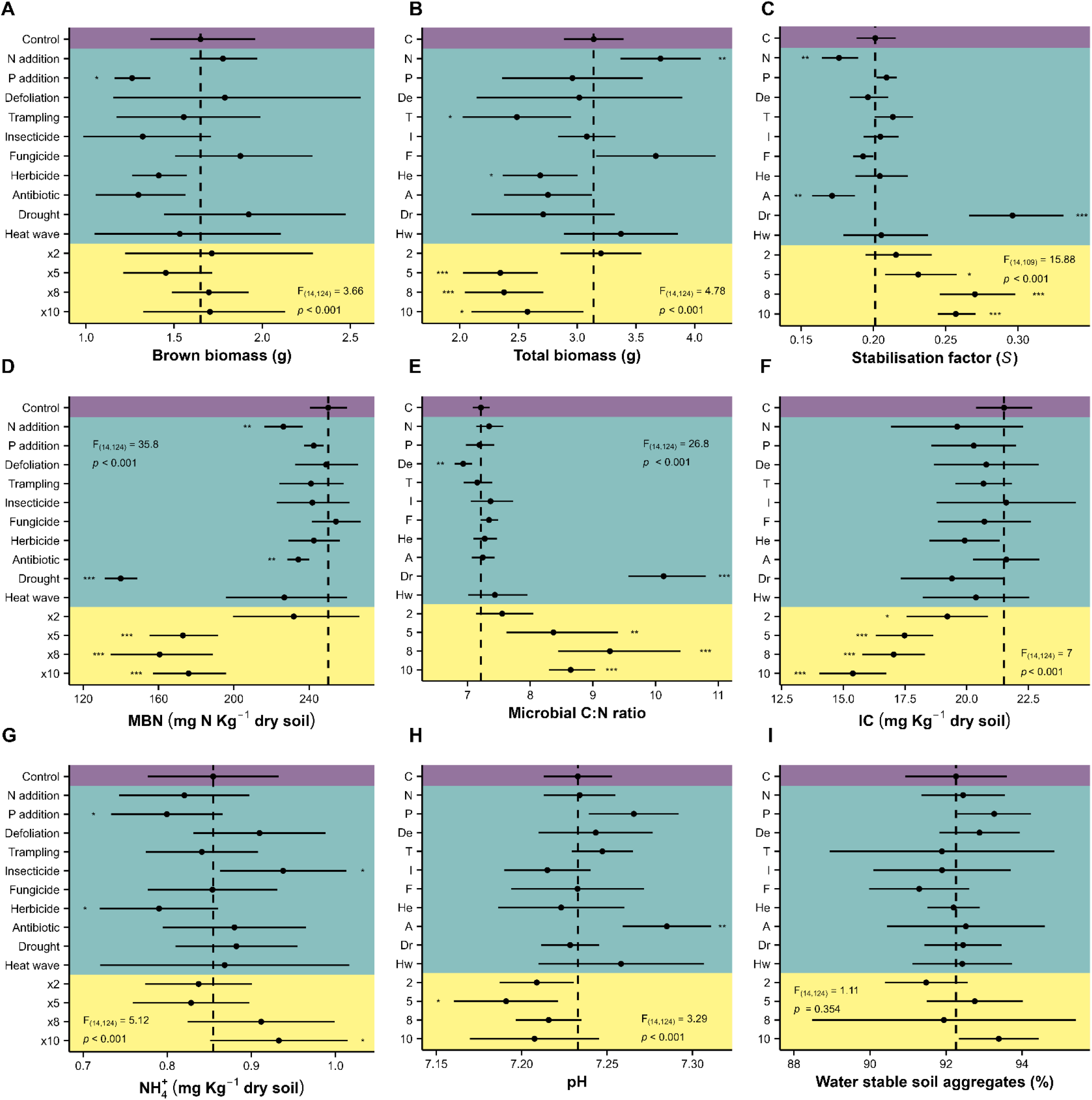
Effects of individual and combined perturbations on the different response variables. MBN = microbial biomass nitrogen, IC = soil inorganic carbon. Estimated marginal means of the model plus 95% interval confidence are shown. Significance of the effect is shown as *: p < 0.05, **: p < 0.01, ***: p < 0.001.

**Fig. S4:**
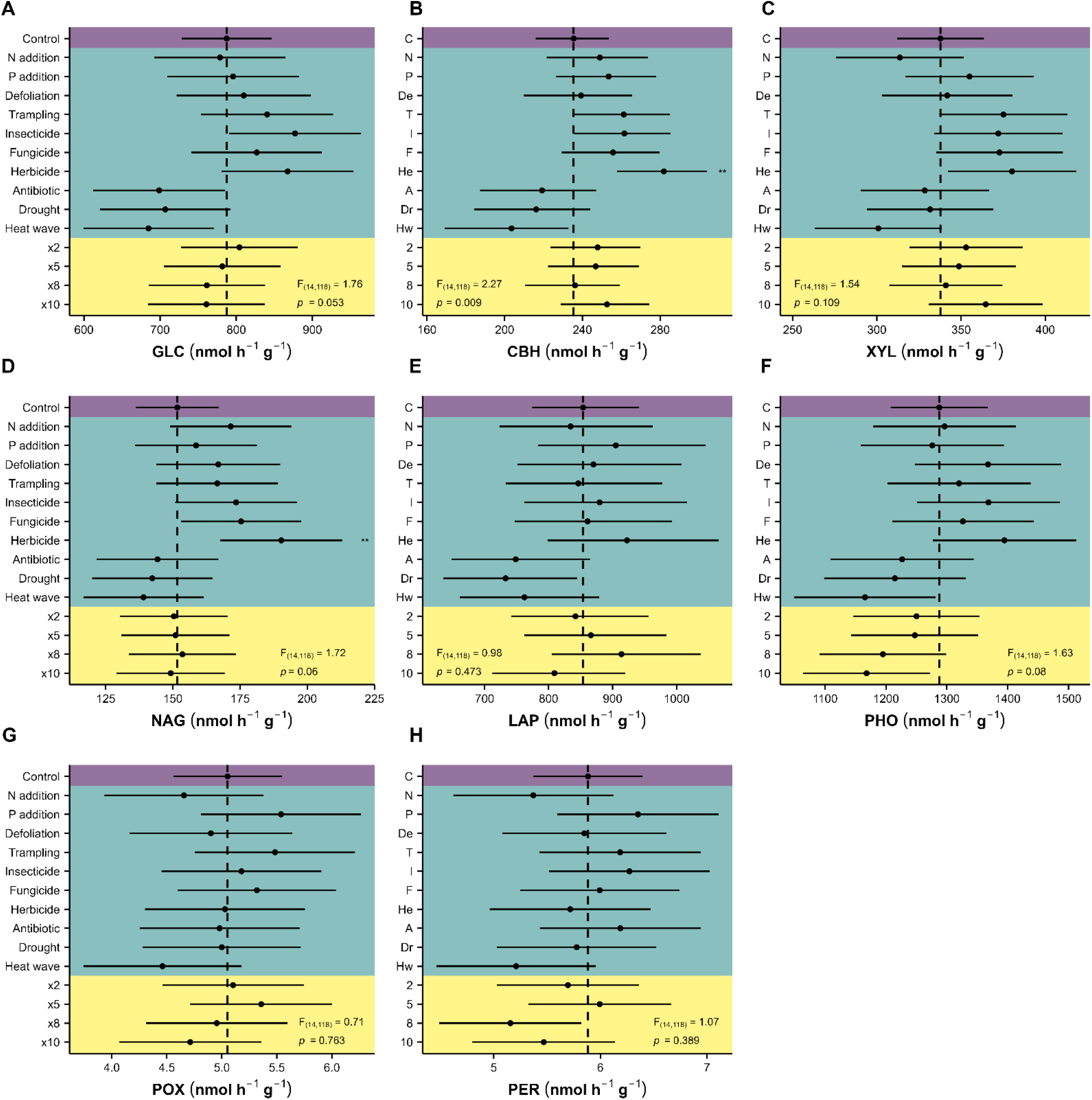
Effects of individual and combined perturbations in the different soil enzymatic activities. GLC = *β*-glucosidase, CBH = cellobiohydrolase, XYL = xylosidase, NAG = *N*-acetylglucosaminidase, LAP = leucine aminopeptidase, PHO = acid phosphatase, POX = phenoloxidase, PER = peroxidase. Estimated marginal means of the model plus 95% interval confidence are shown. Significance of the effect is shown as *: p < 0.05, **: p < 0.01, ***: p < 0.001.

**Fig. S5:**
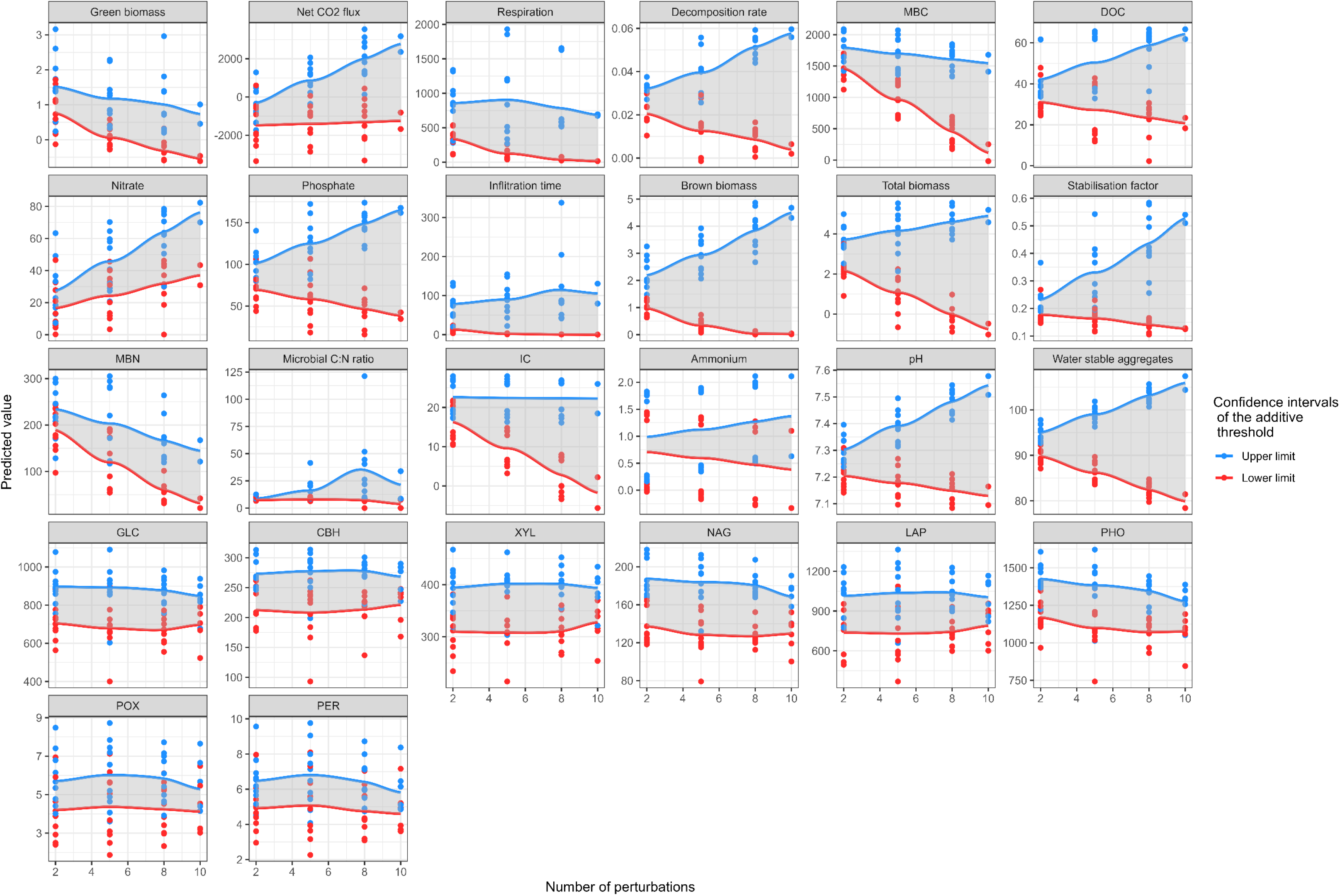
Confidence intervals of the predictions of the additive model for all measured response variables. Lines represent smoothed trends of the confidence intervals with increasing number of perturbations. Shaded area represents the width of the confidence intervals (distance from upper limit to lower limit). Plant biomass (green, brown and total), in g. Net CO_2_ flux and respiration, in mg CO_2_ m^-2^ h^-1^. Microbial biomass and soil nutrients, all in mg N or C Kg^-1^ dry soil: MBC = microbial biomass carbon, MBN = microbial biomass nitrogen, DOC = dissolved organic carbon, IC = soil inorganic carbon. Infiltration time, in s. Water stable aggregates, in %. All enzymes in nmol h^-1^ g^-1^: GLC = β-glucosidase, CBH = cellobiohydrolase, XYL = xylosidase, NAG = N-acetylglucosaminidase, LAP = leucine aminopeptidase, PHO = acid phosphatase, POX = phenoloxidase, PER = peroxidase. Decomposition rate (*k*), Stabilisation factor (*S*), Microbial C:N ratio and pH, unitless.

**Fig. S6:**
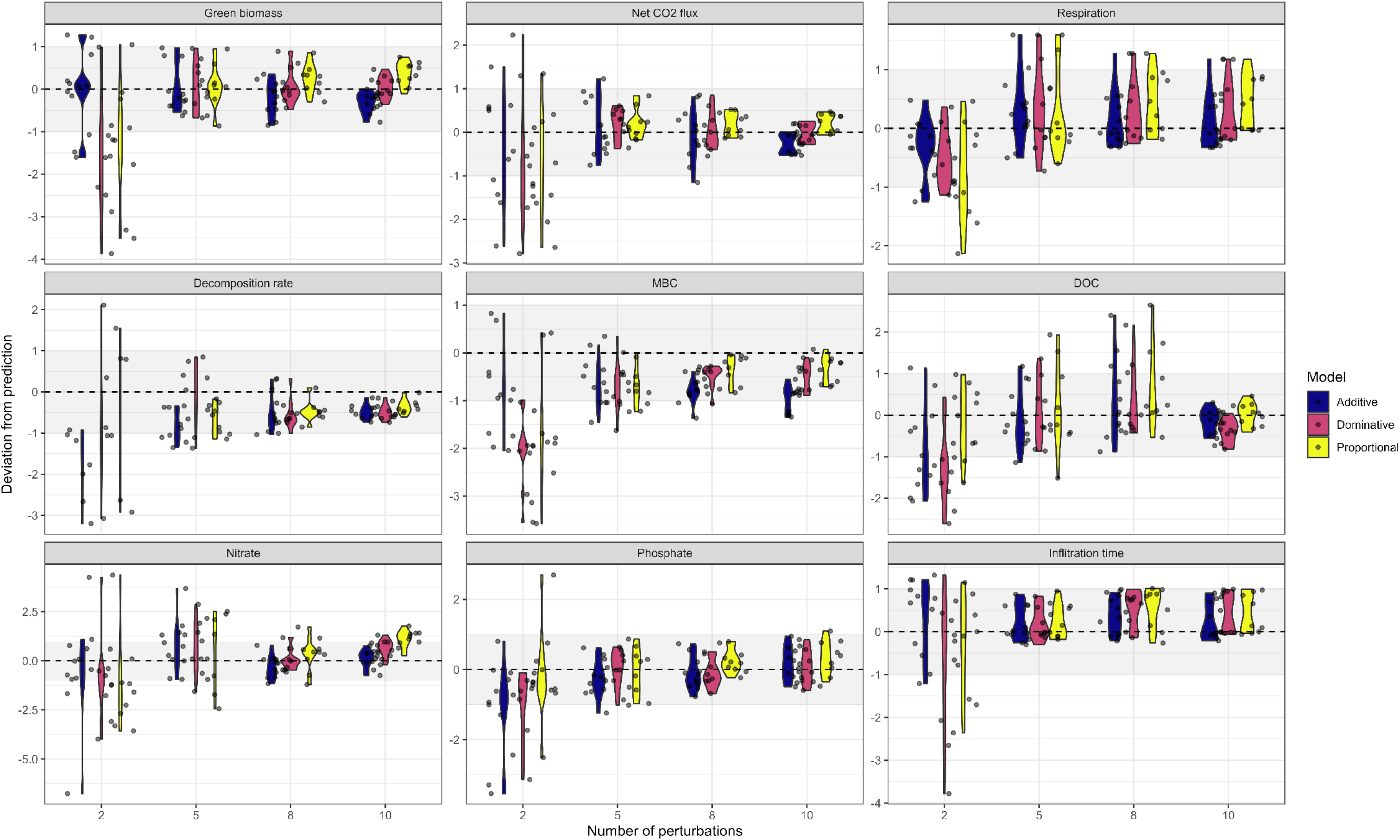
Deviation of observed effects of multiple perturbations from the predictions of three different null models: additive, dominative and proportional. Predictions were made based on the response to single perturbations. Grey shaded areas indicate the 95% confidence interval of the prediction, which was scaled between 1 and −1. MBC = microbial biomass carbon, DOC = dissolved organic carbon.

**Fig. S7:**
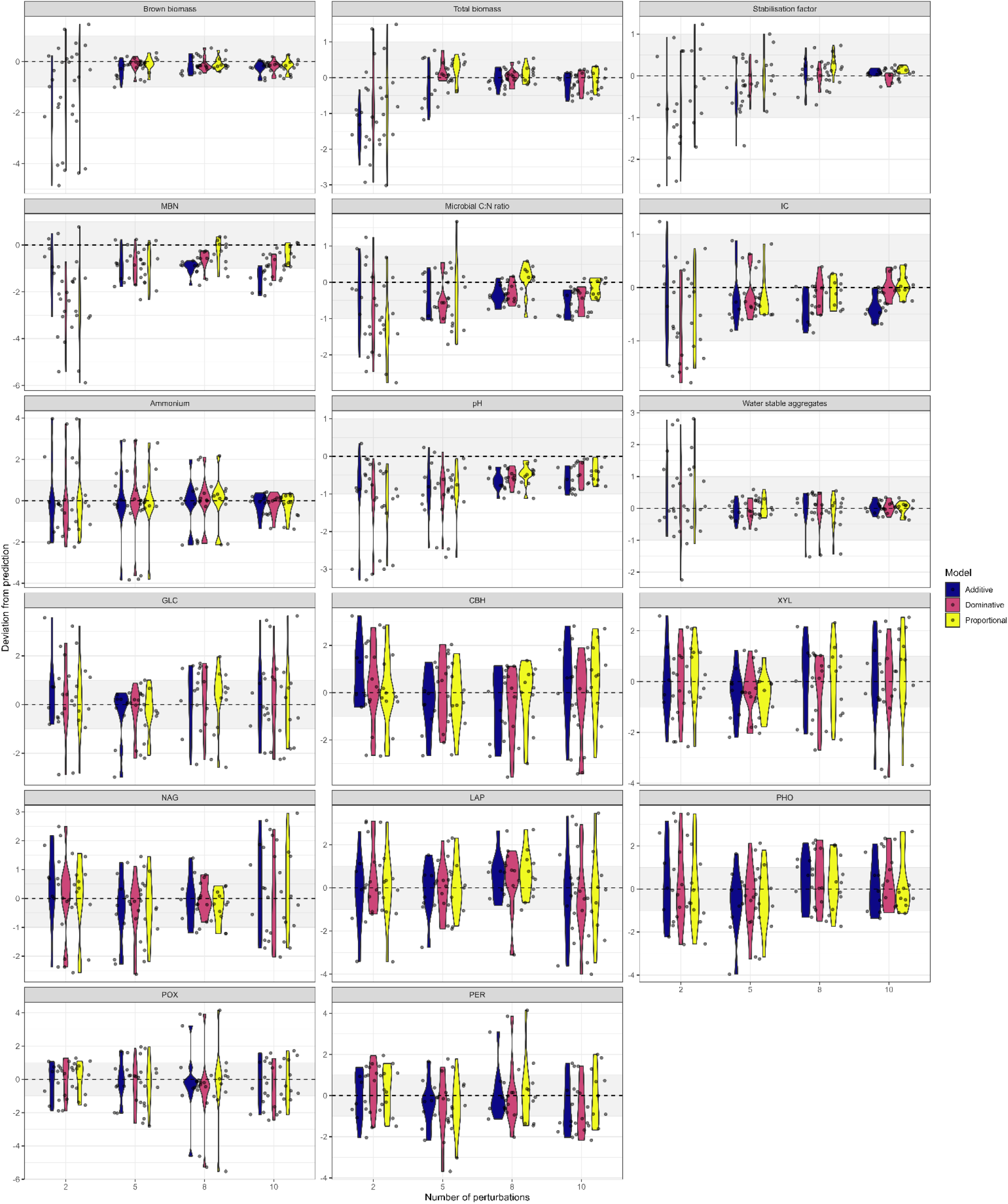
Deviation of observed effect of multiple perturbations from the predictions of three different null models: additive, dominative or proportional. Predictions were made using the response to single perturbations. Grey shaded areas indicate the 95% confidence interval of the prediction, which was scaled between 1 and −1. MBN = microbial biomass nitrogen, IC = inorganic carbon, GLC = β-glucosidase, CBH = cellobiohydrolase, XYL = xylosidase, NAG = N-acetylglucosaminidase, LAP = leucine aminopeptidase, PHO = acid phosphatase, POX = phenoloxidase, PER = peroxidase

## Notes

### Competing Interest Statement

The authors have declared no competing interest.

